# Fine-scale ecological biomonitoring in a large, complex agriculturally impacted watershed via eDNA metabarcoding

**DOI:** 10.1101/2025.11.24.690238

**Authors:** Bráulio S. M. L Silva, Andrew C. Riley, Emilia Craiovan, Michael Wright, Katherine Watson, David R. Lapen, Mehrdad Hajibabaei

## Abstract

DNA-based approaches utilizing high-throughput sequencing (HTS) (e.g. DNA metabarcoding) have revolutionized ecological biomonitoring by providing higher sample throughput, greater reproducibility, and better cost-benefits compared to traditional morphology-based bioassessment studies. Here, we utilized DNA metabarcoding in a watershed in Ontario (Canada) dominated by agricultural land uses. Our aim is to understand patterns of biodiversity in benthic taxa from data generated and inferred at various taxonomic scales and to compare these findings with over a decade of traditional morphological data. We sampled 18 watercourses during summer and fall 2023, spanning a forested-to-agricultural land-use gradient. We found significant differences between metabarcoding and historical morphology data where DNA provided more richness values at both the species (p = 2×10^-5^) and order (p = 0.008) levels. Whereas the morphology dataset contained many unresolved taxa, DNA metabarcoding captured a broader taxonomic breadth with diverse genetic profiles among taxa. Non-metric multidimensional scaling (NMDS) analyses on DNA metabarcoding data produced tighter clusters, more precise separation by land use, and greater consistency across taxonomic scales. Both urban context and land use had significant associations with metabarcoding patterns observed, with differences being strongest between agriculturally-dominated and primarily forested sites (median R² ≈ 0.08-0.11). We also found strong, consistent environmental signals linked to agricultural settings, such as water conductivity and turbidity, and pH. Altogether, our DNA-based results demonstrate the differences in community composition among different land uses in this watershed. Importantly, our work highlights the need for more taxonomic resolution (obtained through DNA analysis) to decipher community changes linked to anthropogenic and environmental drivers, as morphological data alone may lack the precision needed to capture these patterns.

## Introduction

Agricultural extensification and urbanization are significant drivers of reductions in biodiversity globally (Ellis et al., 2013; Ramankutty et al., 2018). Agriculture is the most extensive form of impacting land use, occupying more than 37% of the Earth’s terrestrial surface, while urban areas are expanding at unprecedented rates (Ellis et al., 2013; Klein Goldewijk et al., 2017; Almond et al., 2020; Schiavina et al., 2022; Zhang et al., 2025). These expanding land uses are of particular concern for freshwater ecosystems, as both agriculture and urban development can have profound physical and chemical effects on such systems via degrading water quality and hydro-physical habitats (Dudgeon et al., 2006; Geist, 2011; Saxena, 2025; Mishra, 2025).

The physical consequences resulting from environmental alterations are well documented. Removal of riparian vegetation, stream channelization, and modified flow regimes lead to habitat simplification and loss. For example, elimination of vegetation along riparian buffers can increase stream temperatures, reduce protection from ultraviolet radiation, and make watercourses more vulnerable to erosion, nutrient inputs, and sediment loads (Osborne and Kovacic, 1993; Kelly et al., 2003; Sweeney et al., 2004; Cole et al., 2020). Such changes not only compromise habitat quality but also disrupt the ecological features that maintain freshwater biodiversity.

In Ontario, Canada, agricultural land drainage has contributed significantly to wetland loss, with some counties experiencing reductions of nearly 50% of their original wetland cover (Spaling, 1995; Birch et al., 2022; Penfound and Vaz, 2022). A key driver of this loss has been the widespread installation of tile (artificial subsurface) drainage systems, which artificially drain millions of hectares of farmland and remain one of the most common practices in the region (Sunohara et al., 2014, 2015; Que et al., 2015; Controlled Tile Drainage in Ontario: Cost-Benefit Analysis, 2017). The South Nation River (SN) basin is one of the most significant watersheds in Eastern Ontario, covering an approximate area of 3,900 km². While land use is mixed, it is predominantly agricultural, with widespread field crop production supported by extensive subsurface tile drainage systems (Crabbé et al., 2012; Sunohara et al., 2015; Noteboom et al., 2021).

The SN watershed has been part of a long-term ecological monitoring project by the South Nation Conservation Authority (SNCA), where at least 72 fish species were found to occur across streams in the watershed, including *Cyprinus carpio* (carp), *Sander vitreus* (walleye), *Micropterus dolomieu* (smallmouth bass), and *Esox lucius* (pike) (South Nation Conservation Authority, 2014, 2023, n.d.). Other recent studies have explored and described additional biodiversity in the area, from investigating the dietary and nutritional linkages between tree swallows, riparian insects, and emerging aquatic insects to understanding how environmental factors in this agriculturally dominated area affect fungal diversity (Pham et al., 2024; Rideout et al., 2025).

Given the need to track ecological responses in a standardized way, biomonitoring methods have been widely adopted for assessments in agricultural areas, with benthic macroinvertebrates emerging as a central focus for researchers, environmental monitors, and stewards. Most freshwater benthic macroinvertebrates are insects, but the group also includes crustaceans, gastropods, bivalves, and oligochaetes such as aquatic worms that inhabit the substrates of streams, rivers, and lakes (Bae et al., 2005; Menezes et al., 2010; Hussain, 2012; Tampo et al., 2021; U.S. EPA., 2025). These organisms are important to freshwater ecosystems by contributing to nutrient cycling within aquatic food webs and influencing other ecosystem processes such as microbial production and nutrient cycling (Covich et al., 1999; Schmera et al., 2017). Among them, the larval stages of several insect groups, such as the EPT taxa (Ephemeroptera, Plecoptera, and Trichoptera), are especially valuable for biomonitoring. Representative examples include the flat-headed mayfly (Heptageniidae - Ephemeroptera), green sedge caddisfly (Rhyacophilidae - Trichoptera), and golden stonefly (Perlidae - Plecoptera), which are widely recognized as sensitive indicators of ecosystem condition (Resh, 2008). Their sensitivity is partly explained by their close association with water quality, as most benthic macroinvertebrates exhibit limited dispersal during their aquatic larval stages. Furthermore, the relatively long larval periods of some species, which can range from several months to multiple years, make them responsive to both seasonal and interannual environmental changes (May, 2019).

In Ontario, stream condition is commonly assessed using the Ontario Benthos Biomonitoring Network (OBBN) protocol, which sets a consistent approach for kick-and-sweep collection of benthic macroinvertebrates, morphological identification, and metric-based assessment (e.g., EPT richness, Hilsenhoff Family Biotic Index, and trait or feeding-group composition) (Jones et al., 2007). For years, the OBBN protocol has been used in most biomonitoring studies conducted in the SN watershed and other basins in the Province. However, morphology-based approaches to taxonomic identification are often slow, labor-intensive, and limited to certain groups, which can reduce both efficiency and precision. To address these limitations, our group introduced the concept of “Biomonitoring 2.0”, a framework that empowers biomonitoring with DNA-based taxonomic identification using high-throughput sequencing technologies such as DNA metabarcoding (Baird and Hajibabaei, 2012). This approach generates large volumes of genetic data, allowing for species-level resolution that surpasses conventional morphological methods. The genetic information on community composition is obtained through DNA metabarcoding, which involves extracting and sequencing standard gene regions (e.g. DNA barcodes) from environmental or bulk samples (e.g., water, soil, air, or benthic invertebrate collections). Depending on the study objectives and target organisms, different genetic markers can be used for metabarcoding, including the internal transcribed spacer (ITS) for fungi, plastid genes such as matK and rbcL for plants, and the mitochondrial cytochrome c oxidase subunit I (COI) for animals (Hajibabaei et al., 2016).

Despite the growing adoption of DNA metabarcoding for freshwater biomonitoring, several key questions remain. For instance, one important gap is our limited understanding of how DNA metabarcoding performs in highly impacted regions, where conventional morphology-based identification methods have traditionally been applied. This will allow a direct comparison of biomonitoring outcomes and ultimately help with decision-making for adoption of the new DNA-based approach. There remains a shortage of studies evaluating the applicability, effectiveness, and compatibility of these newer methods in long-term monitoring programs (Hering et al., 2018; Leese et al., 2018; Cordier et al., 2021). Another important consideration is the appropriate level of taxonomic resolution required for biomonitoring analyses, especially when morphology-based approaches fail to detect and assign specimens to finer levels, such as species and genus. Finally, access to finer scale genetic data (e.g. Exact Sequence Variants (ESVs)) will provide a new layer of biological information, which is independent of taxonomic inference and can be used as an additional data layer for biomonitoring (Porter and Hajibabaei, 2020; Riley et al., 2025b). Addressing these questions is critical to evaluating the effectiveness of DNA metabarcoding for freshwater bioassessment. For this purpose, we directly compare one year of DNA metabarcoding data with a decade of traditional morphology-based benthic macroinvertebrate surveys in the South Nation watershed in eastern Ontario, Canada.

## Material and methods

### Sample collection

Kick-net samples were collected from 18 stream sites across the South Nation River watershed (eastern Ontario, Canada) during the summer and fall of 2023 (Figure 1). Site selection was based on differences in physiography to ensure a broad spatial representation of surface water systems in the watershed. Field procedures followed the STREAM Method for Collecting Benthic Macroinvertebrate DNA Samples in Wadeable Streams v2.0 (Maitland et al., 2024). Replicate effort was habitat-specific per the protocol: at riffle/run habitats we collected three independent 3-minute kick-net samples, whereas slow-flow/pool habitats were sampled with one 2-minute sweep. The sweep approach is necessary in areas of weak flow, where insufficient water current would otherwise prevent dislodged organisms from entering the kick-net (Chessman, 1995; Davis et al., 2006).

**Figure 1.**
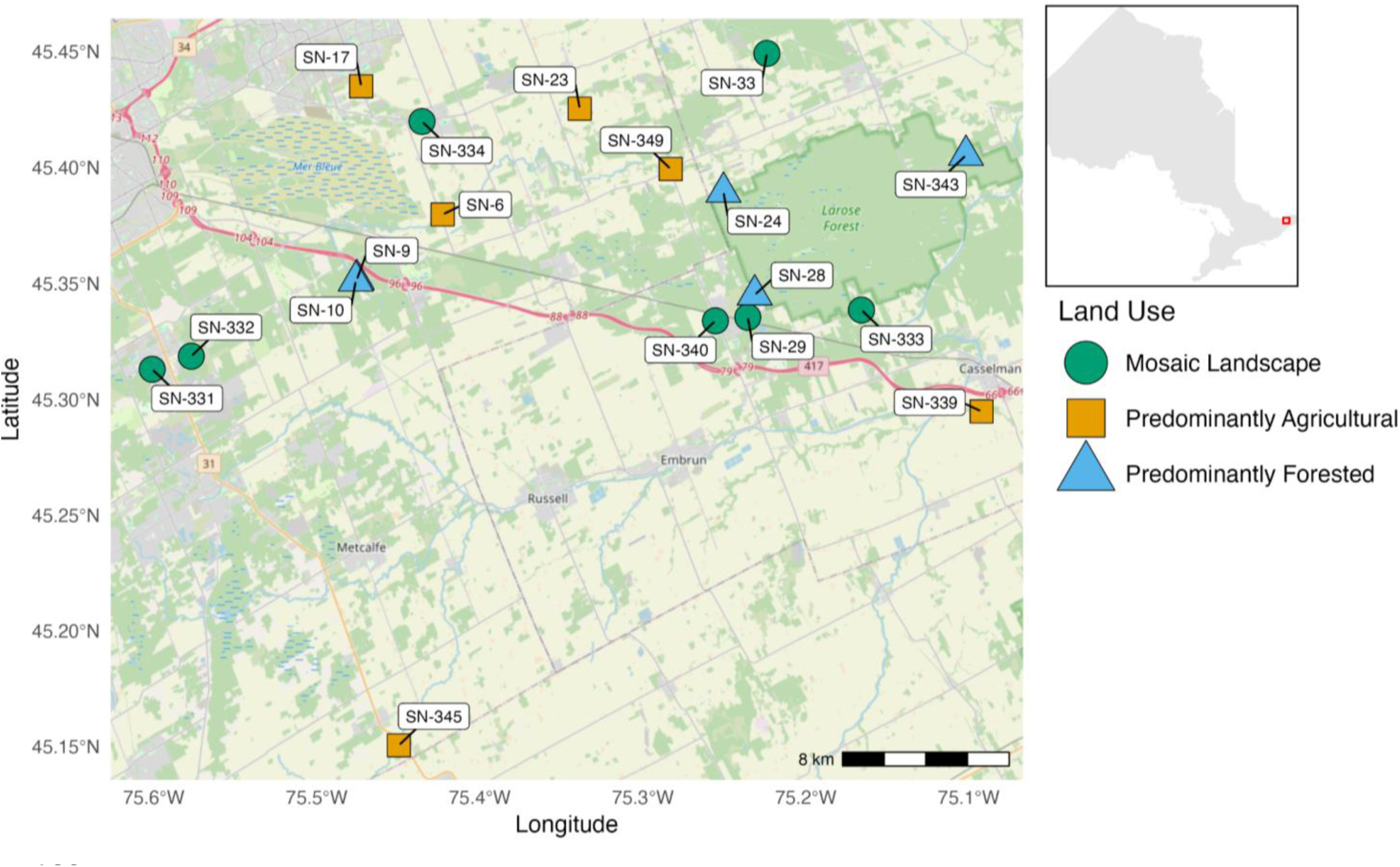
Map of the 18 selected stream sampling sites in the South Nation watershed (eastern Ontario, Canada). Sites are colored and shaped based on the “Land-Use Category”, where: orange square = Predominantly Agricultural (agriculture > 60%), blue triangle = Predominantly Forested (tree cover > 60%), and green circle = Mosaic Landscape (no single category > 60%, with at least two land cover types > 20%). Number of sites (n) for each land-use category: Predominantly Agricultural = 6 (agricultural average: 78.12%), Predominantly Forested = 5 (forest average: 87.26%), and Mosaic Landscape = 7 (agricultural average: 31.70% and forest average: 40.69%). Site codes are labeled for reference, and the inset map shows the regional context within Ontario, Canada. Land-use classifications were derived from spatial land cover analyses to characterize environmental gradients influencing macroinvertebrate communities (see Material and Methods and Table S5).

We used a YSI Pro DSS probe to measure water temperature (°C), pH, specific conductivity (μs/cm), dissolved O₂ (mg/L and %), turbidity (NTU), and oxidation-reduction potential (mV). These metrics were later applied in environmental fitting analyses (envfit) and correlated with community diversity.

Morphological data were collected by South Nation Conservation (SNC; https://www.nation.on.ca) from 2008 to 2022 using the standard OBBN protocol (Table 1). To compare this morphology-based biomonitoring approach to our one-year DNA metabarcoding assay, we accessed SNCA’s long-term (over 10 years of sampling) benthic macroinvertebrate datasets, which integrate morphological and biological assessments as part of a broader stream condition monitoring strategy. For consistency of comparisons between sampling methods, the morphological dataset was also pre-filtered to remove all non-arthropod taxa (e.g., mollusks, annelids).

**Table 1:**
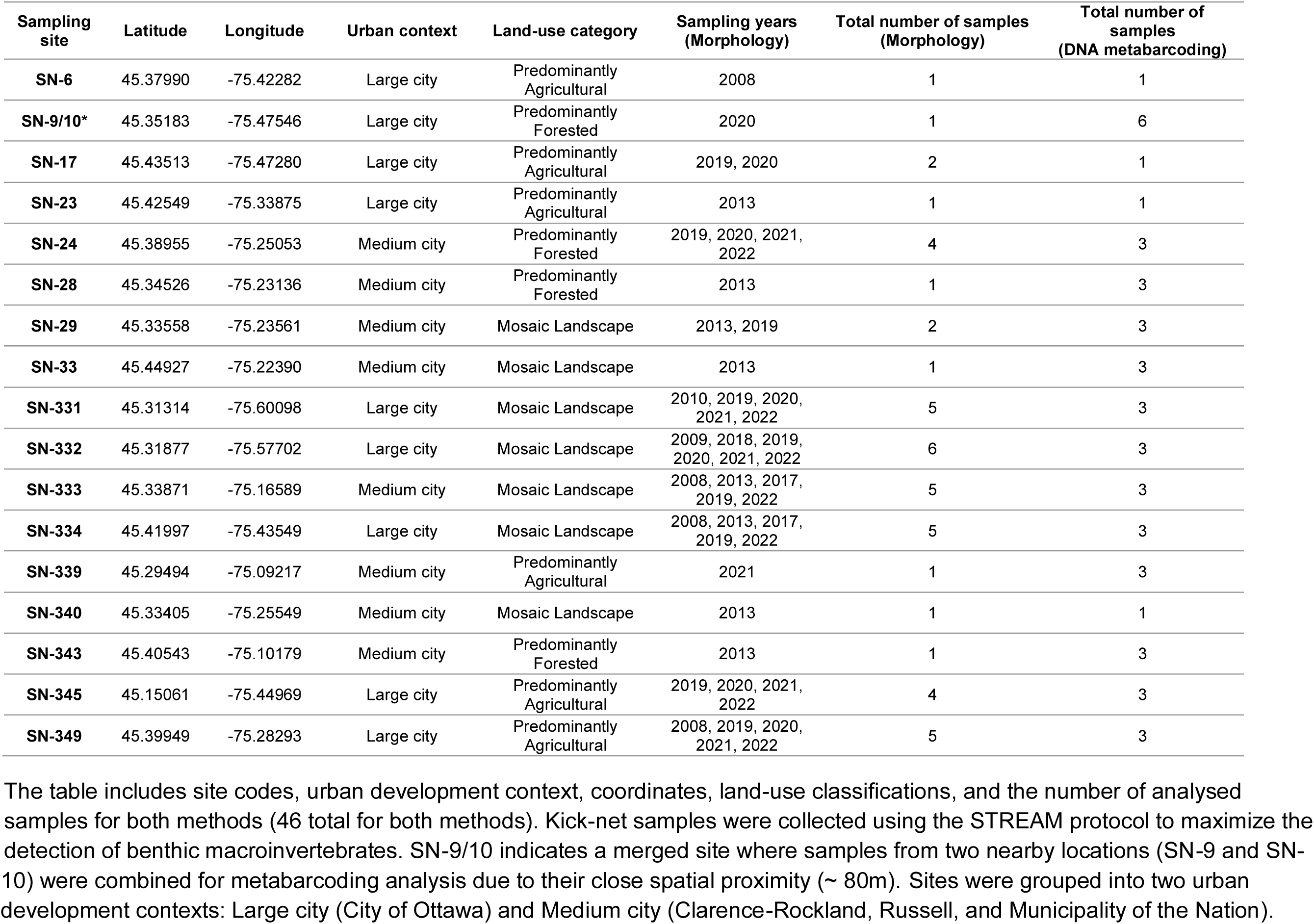
List of sampling sites in the South Nation watershed.

### Land-use categorization

To categorize land use around each sampling site, a “shingled approach” was used. First, the catchment area for each site was identified using a Digital Elevation Model (DEM) obtained from the Ontario Geospatial Data Exchange database (accessed in June 2025). Site coordinates were imported into ArcMap, where 1 km and 2 km buffers were created around each sampling point. These buffers were then used to clip the catchment areas, creating a layer for each site corresponding to each buffer distance. Land use data from the Open Government Portal were clipped to the basin areas using the same 1 km or 2 km buffer, with projections set to NAD_1983_UTM_Zone_18N. The clipped raster was converted into polygons, and the attribute table was used to calculate the area for each land-use category (e.g., agriculture, urban, wetland, forested). A pivot table was created to sum the reclassified areas and determine the percentage of total land area for each site (see Table S5). To investigate the presence of “mosaic” or “transitional” areas, which may represent a gradual shift from minimally impacted environments to more human-influenced or altered landscapes, we established three main land-use categories (1km buffer), defined as: 1) “Predominantly Agricultural: Agriculture > 60%”, 2) “Predominantly Forested: Tree cover > 60%,” and 3) “Mosaic Landscape: No single category > 60%, with at least two categories > 20%,”. A similar approach for categorizing land use at sites in the South Nation watershed was previously employed by Pham et al., (2024).

In addition to land-use categories, we incorporated an urban development context factor representing the degree of urbanization at each site. This variable was treated as a two-level categorical factor (large city vs medium city), complementing the quantitative land-use metrics derived from land-cover data. Together, these descriptors allowed us to test whether community composition patterns were influenced by both local land-use characteristics and broader urbanization context, ensuring that observed patterns were not solely attributable to either factor alone.

### DNA metabarcoding and taxonomic assignments

Macroinvertebrate bulk samples were processed following the STREAM v2.0 benthic metabarcoding protocol (Maitland et al., 2024). In summary, samples were homogenized using a sterile blender, and a subsample of the homogenate was used for DNA extraction with a modified DNeasy PowerSoil Pro Kit (QIAGEN) protocol to increase DNA yield and inhibitor removal. To enhance species detectability and improve coverage across the COI gene, we used three primer sets targeting partially overlapping regions: COI-BR5, COI-F230R, and COI-MLJG. The BR5 region was amplified using the B_F primer (5′–CCIGAYATRGCITTYICG–3′; Hajibabaei et al., 2012) and ArR5 (5′–GTRATIGCICCIGCIARIACIGG–3′; Gibson et al., 2014). The F230R region was amplified with LCO1490 (5′–GGTCACAAATCATAAAGATATTGG–3′; Folmer et al., 1994) and 230_R (5′–CTTATRTTRTTTATICGIGGRAAIGC–3′; Gibson et al., 2015). The third primer set targeted the MLep region using mlCOlintF (ML-F) (5′–GGWACWGGWTGAACWGTWTAYCCYCC–3′; Leray et al., 2013) and jgHCO2198 (JG-R) (5′–TAIACYTCIGGRTGICCRAARAAYCA–3′; Geller et al., 2013). After two rounds of PCR amplification, samples were quantified, normalized, pooled, and indexed with a Nextera XT Index Kit v2. The final library was purified using AMPure XP magnetic beads and quantified using a Qubit fluorometer. Dual-indexed libraries were sequenced on an Illumina MiSeq platform (2×300 bp, v3 chemistry kit). Negative controls were included during both DNA extraction and amplification to monitor for contamination.

Raw Illumina reads were processed with the MetaWorks v1.13.0 pipeline to ensure high-confidence, arthropod-focused taxonomic assignments suitable for multi-marker metabarcoding analyses (Porter and Hajibabaei, 2022). Sequences were demultiplexed, primer-trimmed with Cutadapt, and then quality-filtered, dereplicated, and denoised into exact sequence variants (ESVs) using VSEARCH and UNOISE3. Chimera filtering was performed, and taxonomic assignments were made with the RDP Classifier trained on the COI v5.1.0 reference database, which includes curated records from BOLD and GenBank (Wang et al., 2007; Porter, 2017). To refine results, ESVs were filtered using a custom grep search to retain only Arthropoda taxa and remove vertebrate sequences. Additionally, putative pseudogenes with very low Hidden Markov Model (HMM) scores were discarded through HMM profile analysis (Porter and Hajibabaei, 2021). Only species-level identifications with a bootstrap support (sBP) ≥ 0.7 were kept to avoid the presence of false positives. According to Porter (2018), this minimum bootstrap support threshold yields approximately 95% correct taxonomic assignments.

### Statistical analyses

To evaluate whether the β-diversity patterns visualized in the Non-metric Multidimensional Scaling (NMDS) ordinations were statistically supported, we applied two non-parametric multivariate tests: PERMANOVA (Permutational Multivariate Analysis of Variance) and ANOSIM (Analysis of Similarities). The tests were conducted on both DNA metabarcoding and morphological datasets, using presence/absence matrices for the former and abundance data for the latter, following standard practices in DNA-based biomonitoring studies. For metabarcoding, analyses were performed independently for each amplicon (BR5, F230R, and MLJG) across multiple taxonomic levels (ESV, Species, Genus, Family, and Order), with land-use category and urban context as grouping variables. Equivalent tests were run on morphological data to enable method comparisons and assess cross-consistency.

All analyses were carried out in R (v4.4.2; R Core Team, 2024). Bray-Curtis dissimilarity matrices were calculated following Riley (2025a) and using the vegan package (v2.6-6) (Oksanen et al., 2024), which also provided functions for NMDS ordination (metaMDS()), PERMANOVA (adonis2()), and ANOSIM (anosim()). Data were manipulated using the packages “dplyr” (v1.1.4) and “tidyr” (v1.3.1), and all figures were generated with “ggplot2” (v3.5.1) (Wickham et al., 2014a, 2014b; Wickham, 2016). The “stats” package was used to apply Benjamini-Hochberg FDR corrections to adjust p-values for multiple testing.

## Results

In total, 79297 specimens were collected but only 5420 specimens of Insecta and 3289 of Malacostraca were morphologically identified to species level through the OBBN method since 2008. Although OBBN provides a standardized and widely adopted framework for benthic macroinvertebrate monitoring, its reliance on morphology-based identification can limit taxonomic resolution. Certain groups were difficult to identify morphologically, resulting in a high proportion of individuals classified only as “Unidentified” at either the genus or species level. At the family level, unidentified individuals were rare overall (0.5% across all years; 413 of 79,297 - Table S1), with modest peaks in 2008 (10.3%) and 2018 (9.3%), but absent in most other years. At the genus level, unidentified records were extremely high in the early years (92.7-100% between 2008 and 2013) but decreased markedly in later surveys, averaging 10-19% between 2017 and 2022. At the species level, unresolved taxa dominated the dataset in all years, comprising 77.9-100% of individuals annually and over 90% in most years. Overall, taxonomic resolution in the morphology dataset was consistently identified at higher levels (class, order, family) and improved substantially at the genus level in recent years, while species-level identifications remained largely unresolved across the entire monitoring period.

The DNA metabarcoding results of benthic macroinvertebrates revealed high taxonomic richness and variation among sites (Table S2). At the species level, we detected a total of 282 unique taxa, with site-level richness ranging from 19 to 78 species (mean = 53). Across all primers, 23.4% of arthropod ESVs (1,356/5,799) achieved a confident binomial nomenclature for species-level assignments (sBP ≥ 0.7), with rates of 33.6% for F230R (586/1,745), 21.5% for BR5 (387/1,796), and 17.0% for MLJG (383/2,258). Richness patterns were consistent at higher taxonomic levels, with 220 genera (25-79 per site, mean = 54), 111 families (14-47 per site, mean = 28), and 23 orders (9-18 per site, mean = 13) identified. Species distributions across sites were highly uneven: out of the 282 species detected, 123 (43.6%) were restricted to a single site, whereas only 21 species (7.4%) were widespread, occurring in 10 or more sites (Table S2). This strong skew toward rare, locally restricted taxa shows the dominance of specialists in the sampled area and likely contributes substantially to the β-diversity across the South Nation watershed.

We also found differences in richness among sites that aligned with both land use and sampling effort (Table 1). For instance, SN-10 (forested, 6 DNA samples) emerged as the most diverse location, with 78 species, 79 genera, 41 families, and 15 orders, followed closely by SN-24 (forested, 3 DNA samples; 72 species, 65 genera, 32 families, 13 orders) and SN-343 (forested, 3 DNA samples; 71 species, 68 genera, 33 families, 12 orders). These forested sites represent biodiversity-rich areas within the watershed and suggest that intact riparian conditions favor the maintenance of diverse macroinvertebrate assemblages. In contrast, sites such as SN-23 (agricultural, 1 DNA sample; 19 species, 25 genera, 17 families, 11 orders), SN-340 (mosaic, 1 DNA sample; 21 species, 29 genera, 14 families, 9 orders), and SN-17 (agricultural, 1 DNA sample; 29 species, 31 genera, 21 families, 10 orders) exhibited the lowest richness values, reflecting the combined effects of stronger anthropogenic pressures and lower sampling intensity (slow flow/pool habitats) (Table S2). Additionally, by running a cross-level richness correlation analysis (also known as “taxonomic sufficiency assessment”), we found a strong positive correlation between genus- and species-level richness (Pearson’s r = 0.97, p < 1 × 10⁻¹⁰, and Spearman’s ρ = 0.96, p < 1 × 10⁻⁹), indicating that genus-level patterns capture nearly all of the diversity signal in our dataset. This is particularly valuable in biomonitoring applications, where incomplete DNA reference databases or morphological limitations often constrain species-level resolution. Similar conclusions have been drawn in other aquatic biomonitoring studies, where genus-level identifications have been shown to capture most ecological patterns observed at the species level (Lenat and Resh, 2001; Jones, 2008).

### Alpha-diversity analysis

In a comparative analysis between our DNA metabarcoding dataset and the historical morphology dataset, we found a total of 303 species confidently assigned to 23 orders (Figure 2A). Of these, 261 species (86.1%) were detected only by DNA metabarcoding, 22 (7.3%) appeared only in the morphology records, and 20 (6.6%) were detected by both approaches. Species richness was concentrated in a few orders as follows: Diptera (90 species - 29.7% of the total; 98% DNA-only, 2% both), Trichoptera (41 - 13.5%; 90% DNA-only, 7% morphology-only, 2% both), Coleoptera (35 - 11.6%; 57% DNA-only, 23% morphology-only, 20% both), and Ephemeroptera (32 - 10.6%; 75% DNA-only, 16% morphology-only, 9% both). Additional well-represented groups included Odonata (17 - 82% DNA-only, 18% both) and Plecoptera (15 - 100% DNA-only). Morphology-only detections were uncommon and concentrated primarily in Coleoptera (8 species), Ephemeroptera (5), Hemiptera (4), and Trichoptera (3), whereas cross-method overlap was greatest in Coleoptera (7 shared species), Ephemeroptera (3), and Odonata (3). Additionally, 62 species (20.5%) of all species were detected exclusively by DNA metabarcoding, spanning 15 orders. These orders are: Plecoptera (15 - 5.0%), Lepidoptera (8 - 2.6%), Hymenoptera (6 - 2.0%), Cyclopoida (5 - 1.7%), Diplostraca (5 - 1.7%), Trombidiformes (4 - 1.3%), Podocopida (4 - 1.3%), Megaloptera (4 - 1.3%), Isopoda (3 - 1.0%), Araneae (2 - 0.7%), Entomobryomorpha (2 - 0.7%), and single-species occurrences in Mesostigmata, Lithobiomorpha, Psocoptera, and Sarcoptiformes (0.3% each).

**Figure 2:**
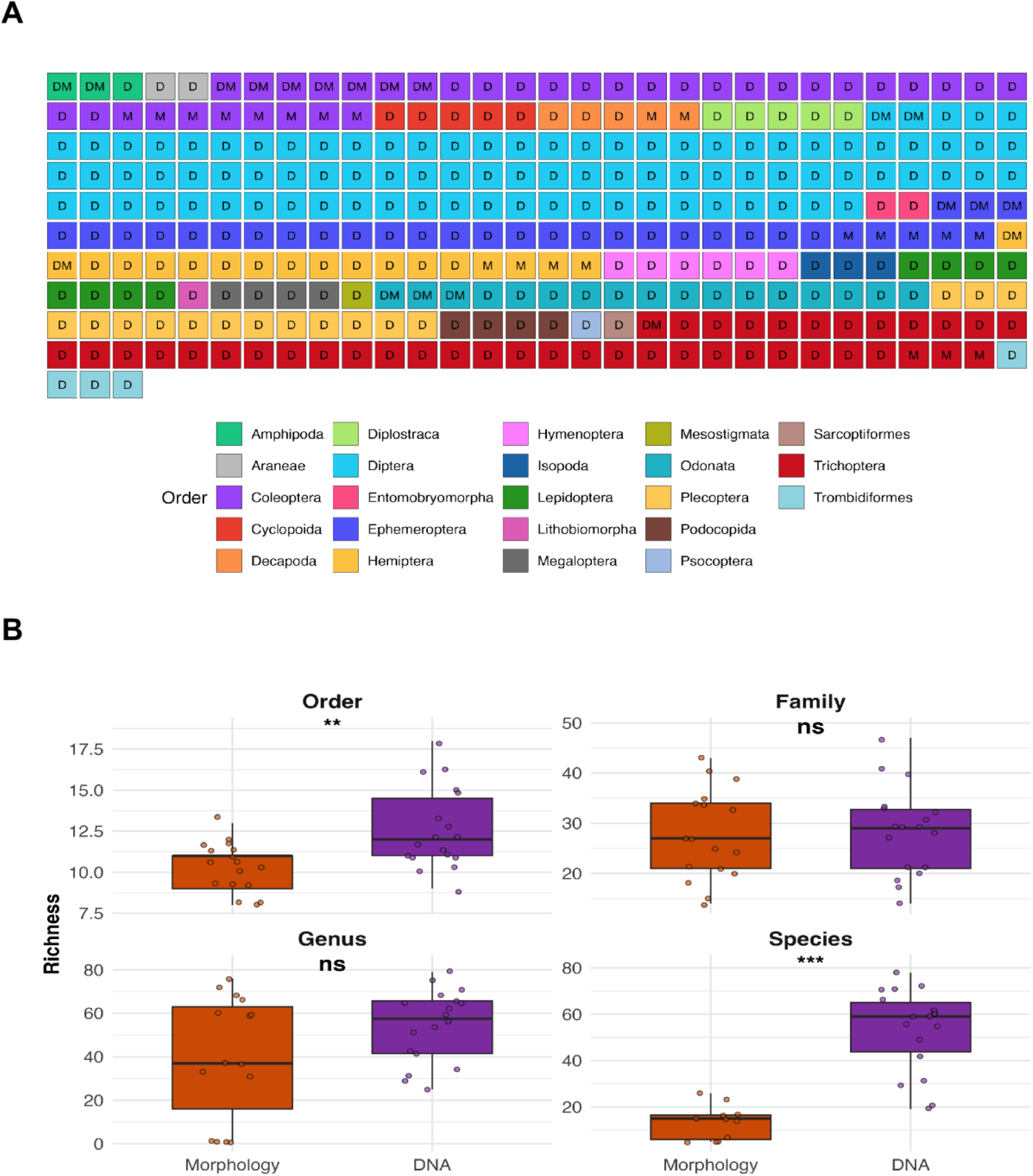
Taxon richness across taxonomic levels: DNA metabarcoding vs. morphology. (A) Species grid by order. Each square represents one species. Tile color denotes the taxonomic order (see legend) and the letter inside indicates the detection method, where: D = DNA metabarcoding only, M = morphology only, DM = both. Orders are shown with a fixed color palette and arranged alphabetically. (B) Site-level richness (number of taxa) by method across taxonomic levels (order, family, genus, species). Each point represents a sampling site. Wilcoxon signed-rank tests were used to assess statistical differences (p < 0.05: *, p < 0.01:**, p < 0.001: ***, ns: not significant). At the species (p-value = 0.00002) and order level (p-value = 0.008), DNA metabarcoding recovered significantly more taxa than morphology, while genus-and family-level richness showed no significant differences.

Across sites, DNA metabarcoding showed higher median richness than morphology at the species and order levels (species p = 2×10⁻⁵; order p = 0.008; Figure 2B). At the species level, medians were 59 (DNA) vs 15 (morphology), indicating a strong DNA advantage consistent with the detection of cryptic or morphologically indistinguishable taxa. At the genus level, medians were 57 for DNA and 37 for morphology, but the difference was not significant (p = 0.261), likely reflecting a larger shared genus pool (86 shared; 30.8%). At the family level, methods were similar (29 for DNA vs 27 for morphology; p = 0.894), although DNA still recovered more unique families overall. At the order level, DNA also showed a small but significant advantage (12 for DNA vs 11 for morphology; p = 0.008). In sum, our comparative analysis indicates that a single sampling campaign using DNA metabarcoding yields higher site-level richness as compared to several years of morphology-based campaigns. This superiority is chiefly reflected at the species and order ranks, while genus and family counts are broadly comparable between methods.

### Taxonomic resolution across macroinvertebrate orders and families

To better characterize the taxonomic structure and genetic diversity of the recovered species, we performed a comparative treemap analysis based on hierarchical classifications from order to family. This visualization approach provides a comprehensive overview of both the breadth of taxonomic coverage and the internal distribution of taxa within and across macroinvertebrate lineages (Figure 3). In the morphology-based dataset (Figure 3A), specimen counts show a community composition dominated by a few abundant families, particularly Elmidae (Coleoptera - 17,672 specimens), Chironomidae (Diptera - 16,801 specimens), and Hydropsychidae (Trichoptera - 11,585 specimens), which together accounted for a significant portion of the observed abundance (46,058 specimens - 58% of total). Families within Ephemeroptera (e.g., Caenidae, Heptageniidae) were also represented but at lower relative abundances, while many additional families fell into the “Others” category due to their comparatively small contributions. Moreover, several taxa remained classified as “Unidentified”, highlighting the difficulty of resolving all taxonomic assignments through morphology-based approaches.

**Figure 3:**
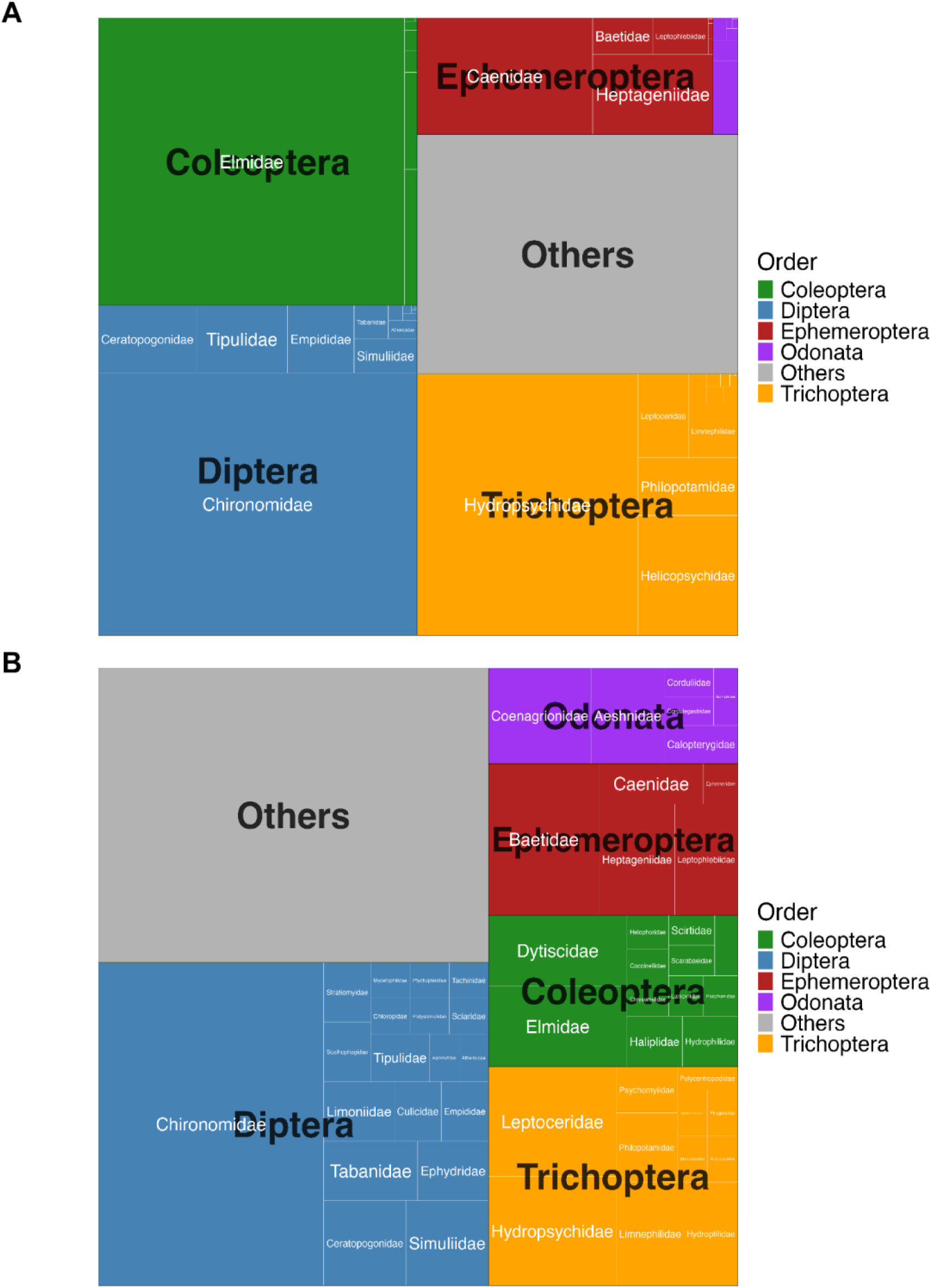
Treemap visualizations of macroinvertebrate community composition based on morphology-based x DNA metabarcoding assessment methods, grouped hierarchically by order and family. Each tile represents a family, with area proportional to (A) specimen abundance from morphological identifications or (B) the number of unique ESVs detected by DNA metabarcoding. (A) Morphology-based identifications emphasize dominant families such as Chironomidae (Diptera), Elmidae (Coleoptera), and Hydropsychidae (Trichoptera). (B) Our DNA metabarcoding assay reveals a broader taxonomic diversity and higher resolution, highlighting additional families such as Dysticidae (Coleoptera) and Aeshnidae (Odonata), and demonstrating the ability of DNA metabarcoding not only to assess community composition but also to infer patterns of genetic diversity beyond simple specimen counts.

In contrast, our DNA metabarcoding results (Figure 3B) revealed not only these dominant insect lineages but also a wider range of taxonomic diversity and higher resolution across families. The order Diptera remained prominent, with Chironomidae occupying the largest tile, reflecting both its ecological ubiquity, taxonomic complexity, and genetic diversity. However, DNA data also highlighted families that were underrepresented or absent in the morphological dataset, including Leptoceridae, Baetidae (Ephemeroptera), and Aeshnidae (Odonata), among others. Additional orders such as Plecoptera, Hemiptera, and Lepidoptera, along with non-insect arthropods (e.g., Amphipoda, Isopoda, Trombidiformes) and crustaceans (e.g., Cladocera, Copepoda) were also detected, expanding the overall taxonomic scope.

### DNA metabarcoding x Morphology-based community compositions

Because our sites had been historically analyzed using morphological data, we opted for site-level comparisons using both historical morphology-based data and newly generated DNA metabarcoding data. The NMDS ordinations revealed consistent differences in macroinvertebrate community composition across land-use categories, with contrasting outcomes between DNA metabarcoding and morphology-based identifications, as well as across DNA amplicons (Figures 4 and 5). Despite representing only a single year of sampling, DNA metabarcoding samples consistently produced tighter within-group clustering and more distinct separations among land-use types for all taxonomic levels analyzed. In contrast, morphology-based ordinations displayed greater within-group variability and frequent overlap among land-use categories, particularly at the family and genus levels (Figure 4). Consistent with the PERMANOVA/ANOSIM tests, land-use category accounted for ∼8.6% of the variation in community composition in the DNA metabarcoding data (Table 2).

**Figure 4:**
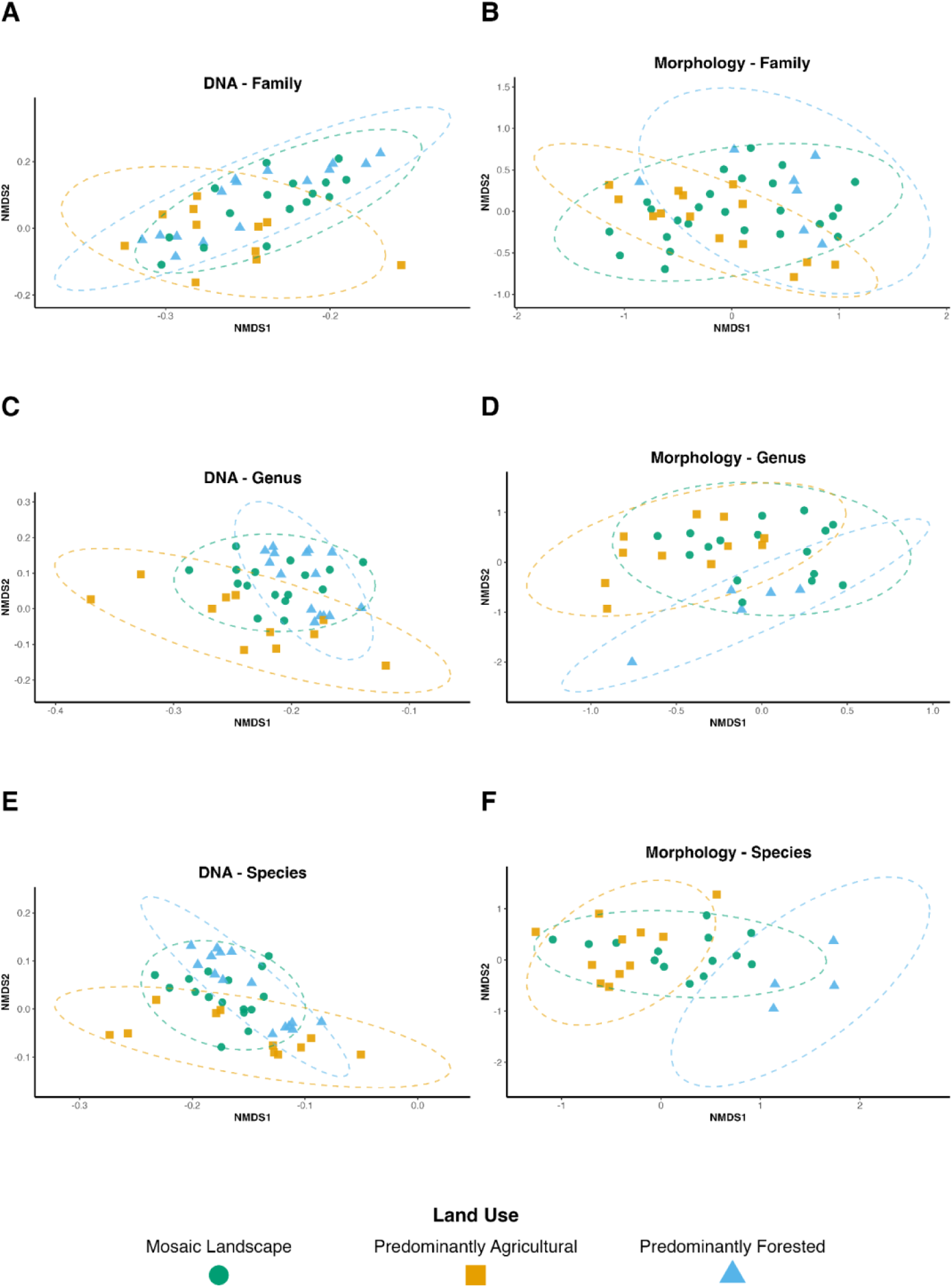
Non-metric multidimensional scaling (NMDS) ordination plots showing macroinvertebrate community composition across land-use types using DNA metabarcoding (left column) and morphology-based identification (right column) at three taxonomic levels: family (A, B), genus (C, D), and species (E, F). Ordinations were computed using Bray-Curtis dissimilarities, based on presence/absence data for DNA (F230R amplicons) and abundance data for morphology. DNA results are based only on one year of sampling (2023), while morphology results represent ten years of sampling. Points represent individual samples, and 95% confidence ellipses group sites by land-use category: mosaic Landscape (green circle), predominantly agricultural (orange square), and predominantly forested (blue triangle). Outlier samples were excluded from the plot based on either low taxonomic resolution (i.e., only a few specimens assigned at the corresponding taxonomic level in the morphology dataset) or aberrant ordination patterns likely resulting from sequencing errors or DNA contamination (see Figure S1). Stress values and dimensions: DNA - family (0.148, k=2), DNA - genus (0.155, k=2), DNA - species (0.146, k=2); Morphology - family (0.164, k=3), Morphology - genus (0.067, k=2), Morphology - species (0.187, k=2).

**Figure 5:**
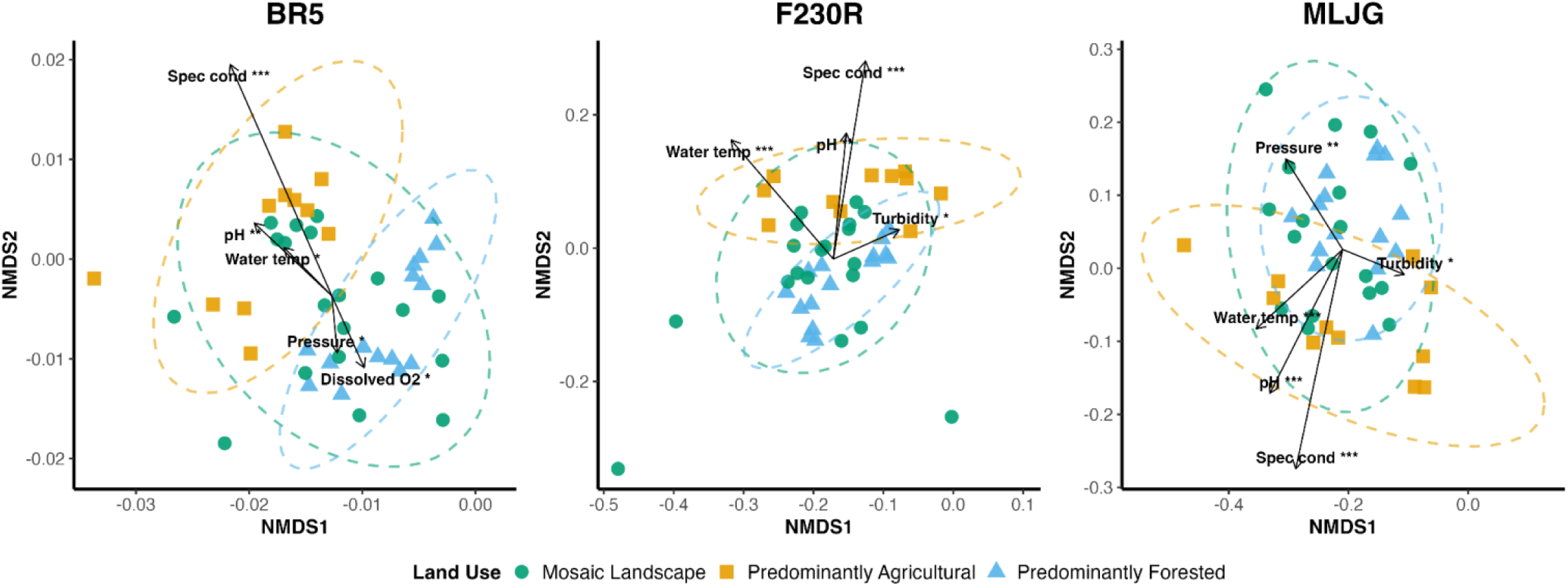
Non-metric multidimensional scaling (NMDS) ordination plots showing macroinvertebrate community composition based on ESV (Exact Sequence Variant) data across land-use categories for three COI amplicons: BR5, F230R, and MLJG. Plots show differences among land-use types, particularly in BR5 and F230R, suggesting that environmental variables such as specific conductivity, dissolved oxygen, and pH play a key role in shaping macroinvertebrate community structure. Points represent samples (replicates included), colored by land-use category: predominantly agricultural (orange square), predominantly forested (blue triangle), and mosaic landscape (green circle). Dashed ellipses indicate 95% confidence intervals around group centroids. Vectors represent significant environmental variables (envfit, *p* < 0.05), scaled relative to the NMDS spread and weighted by the strength of correlation (R²), such that longer arrows indicate stronger associations with community structure. Asterisks indicate levels of statistical significance (*p* < 0.05: *, *p* < 0.01: **, *p* < 0.001: ***) and arrows originate from the centroid of sample distributions in each plot. All values are included in the supplementary material (Table S4). Tested variables: water temperature (“Water temp”), pH, specific conductivity (“Spec cond”), dissolved O₂, dissolved O₂ (%), turbidity, and oxidation-reduction potential (ORP). “Pressure” denotes barometric (atmospheric) pressure and is reported for completeness only. Stress values: 0.095 (BR5), 0.153 (F230R), and 0.146 (MLJG); k=2.

**Table 2:**
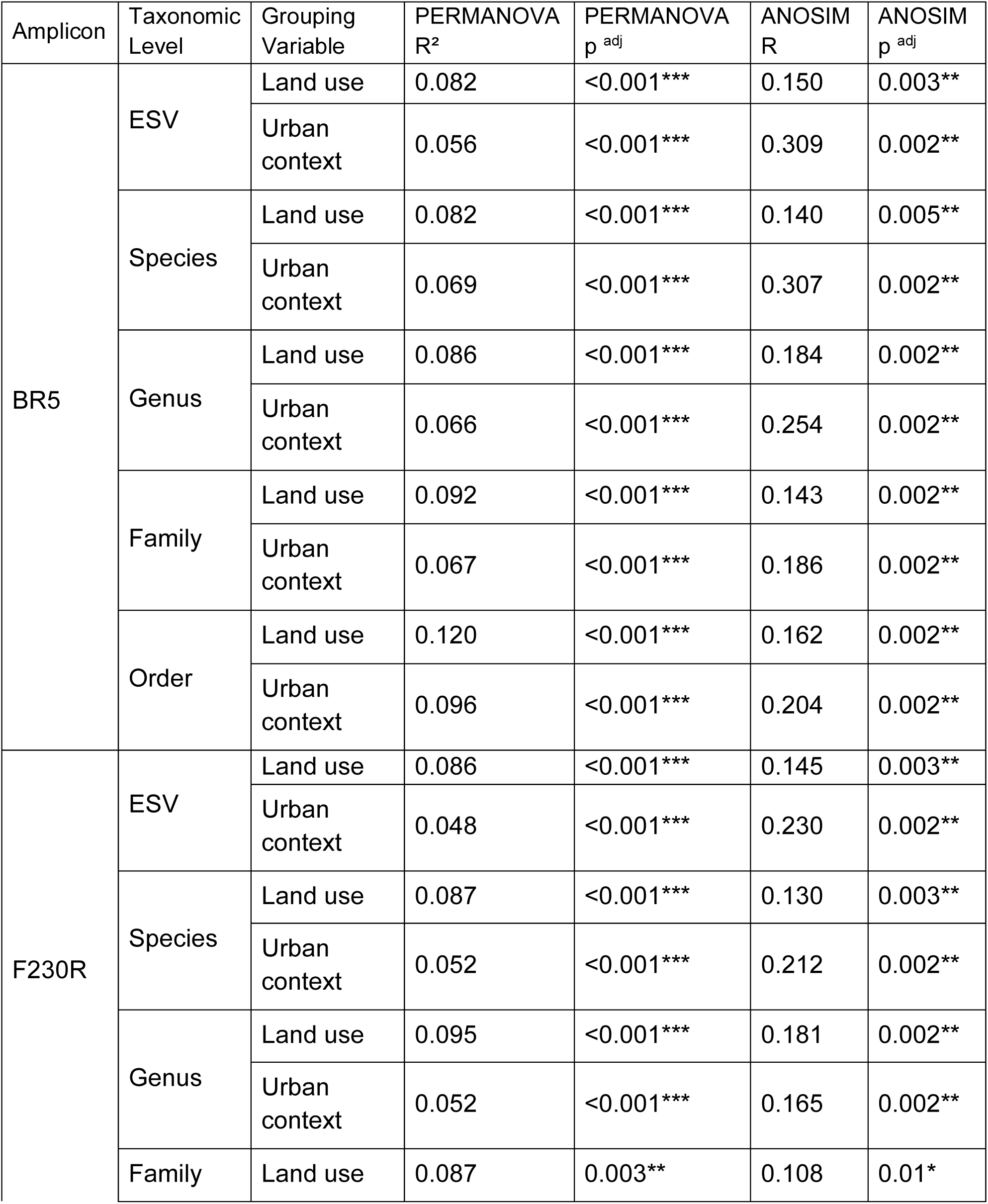
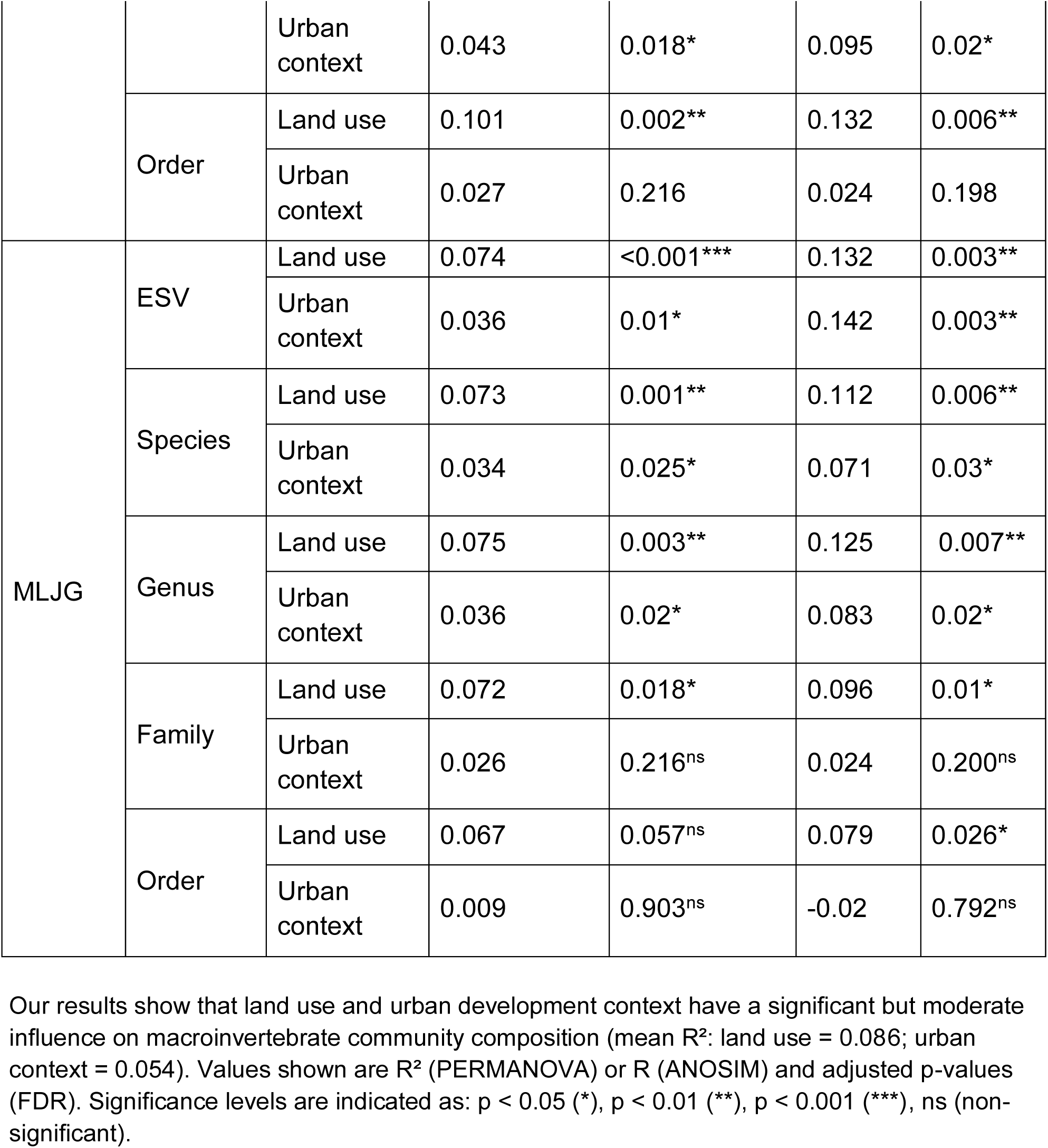
Global multivariate tests of community composition differences across land-use categories and urban development contexts for each amplicon and taxonomic level.

At the species level, morphology achieved its strongest explanatory strength (PERMANOVA R² = 0.178 - p = 0.005; ANOSIM R = 0.359 - p = 0.005 (Table 3)), showing isolating patterns. This pattern was likely driven by a subset of dominant, well-described species that were consistently identified across years and amplified between-group differences, while rarer or cryptic taxa remained unresolved or poorly represented in the morphology data. On the other hand, the DNA metabarcoding data showed consistent, significant community separation at the species level across all three amplicons, with effects stronger for land-use than for urban context (PERMANOVA R² ≈ 0.073–0.087; ANOSIM R ≈ 0.112–0.140; Table 2). Biologically, this indicates that DNA captures a wider share of the species pool (including rare or hard-to-identify taxa), thereby revealing repeatable fine-scale spatial patterns, with urban context differences nested within a stronger local signal. Additionally, DNA metabarcoding leveraged its ability to resolve exact sequence variants (ESVs), capturing subtle within-species turnover and reinforcing the presence of cryptic diversity that were undetectable through morphology (Figure 5).

**Table 3.**
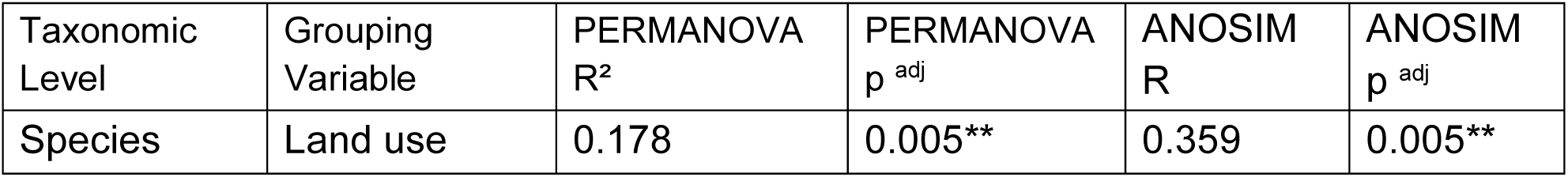
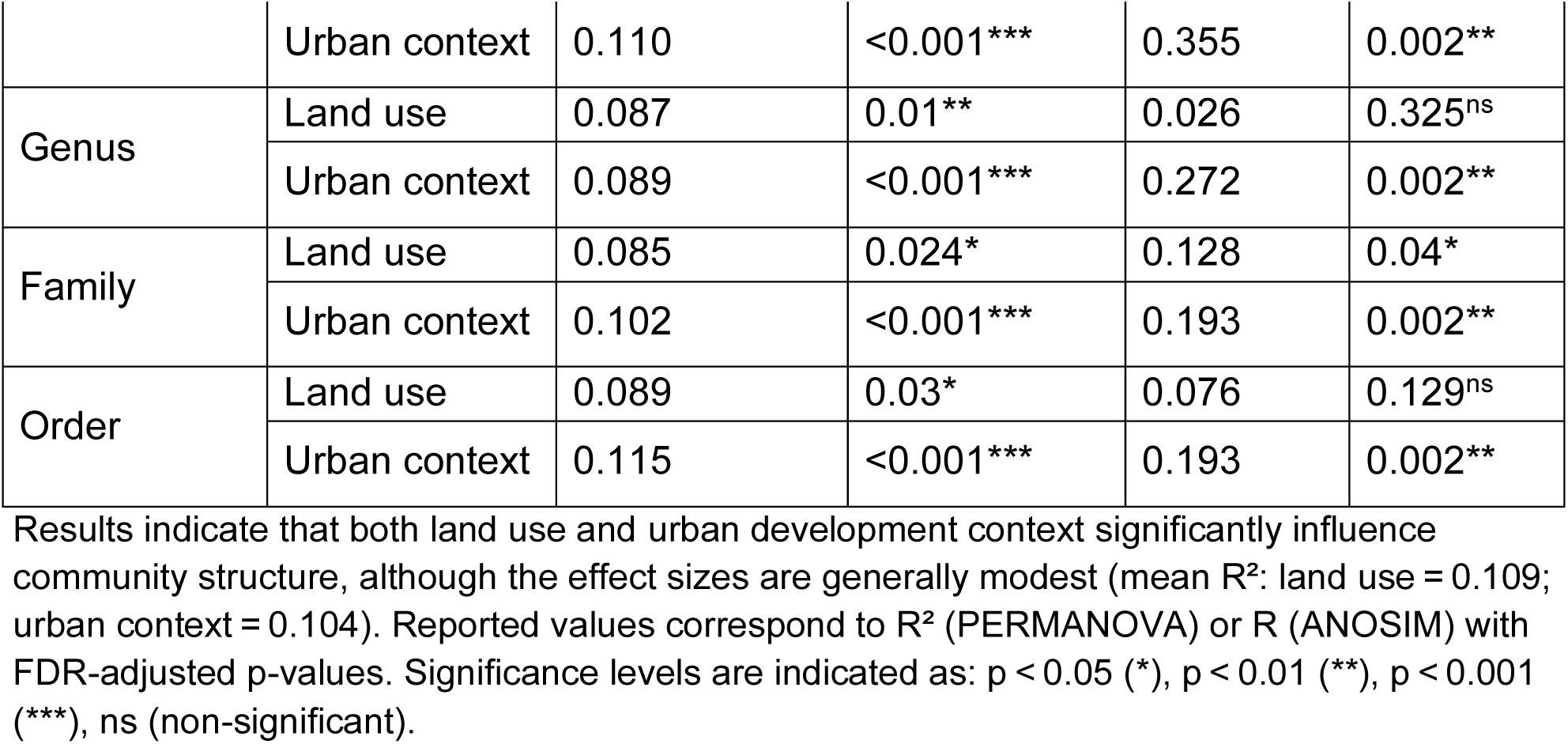
Multivariate tests (PERMANOVA and ANOSIM) assessing differences in morphology-based macroinvertebrate community composition across land-use categories and urban development contexts at multiple taxonomic levels.

Although showing significant results, at low-level resolutions (genus, family, and order), morphology-based patterns became weaker and less consistent. For instance, genus-level analyses showed significant PERMANOVA effects for both land use (R² = 0.087, p = 0.01) and urban context (R² = 0.089, p < 0.001), but ANOSIM failed to detect meaningful land-use differences (R = 0.026, p = 0.325) (Table 3). Similarly, family- and order-level analyses yielded only marginally significant (p=0.04) or non-significant results, suggesting that ecological signals become diluted when taxa are aggregated to higher ranks. A similar result was found for one of the amplicons (MLJG) for DNA metabarcoding (Table 2). Altogether, these results highlight the greater consistency and efficiency of DNA metabarcoding, which achieved comparable or higher resolution with fewer samples and reduced sampling time.

### Environmental Gradients Shaping Community Composition at the ESV Level

Our NMDS results at the genetic level (ESV) combined with envfit statistics showed interesting patterns of community-environment relationships across the COI amplicons (Figure 5; Table S3). In all ordinations, sites were grouped broadly by land use, with agricultural sites generally forming distinct clusters and mosaic and forested sites displaying partial overlap. The envfit analyses revealed that specific conductivity and pH were among the most consistent and strongest correlates across all amplicons. Conductivity explained between 43% and 53% of community variation and was strongly aligned with agricultural sites, reflecting ionic enrichment from fertilizers, manure, and tile drainage, which is typical of intensively managed landscapes, such as farmlands present in the South Nation watershed. Similarly, pH explained 21-38% of the variation, also aligning with agricultural clusters, likely due to nutrient inputs, liming practices, or altered buffering capacity in disturbed streams (Figure 5). These variables highlight the role of physicochemical enrichment as a primary driver of agricultural biocommunity turnover. Turbidity also emerged as a significant correlate (F230R R² = 0.15, p = 0.043 and MLJG R² = 0.18, p = 0.024; Table S3), further supporting the influence of agriculture, where reduced riparian vegetation and soil disturbance contribute to higher sediment loads. In contrast, dissolved oxygen showed a significant association aligned with forested sites (BR5 R² = 0.17, p = 0.032; Table S3). Additionally, water temperature also appeared as a significant correlate in the ordinations (BR5, F230R, MLJG: R² = 0.13, 0.33, 0.29; Table S3), with vectors oriented toward agricultural/mosaic clusters. However, because water temperature in our data mostly reflected the sampling season (Summer: 18.8-25.4 °C; Fall: 7.4-13.6 °C), we also ran a within-season envfit analysis to separate seasonal effects from land-use effects. In these season-stratified analyses, water temperature was not significant for any amplicon (Fall p: 0.20-0.94, Summer p: 0.28-0.94), indicating that, as expected, the pooled temperature signal primarily shows seasonal variation rather than land-use influence.

### Variance partitioning: Land-Use and Urban Context Effects

In the DNA metabarcoding dataset, land use consistently explained more variance (≈ 8.6%) than urban context (≈ 5.4%). These patterns are also confirmed by our variance partitioning analyses (Figure 6). At nearly all taxonomic levels and across all three amplicons, land use explained a larger proportion of variation than urban context, often reaching 8-10% at the family and order levels for MLJG, compared to 0-1% explained by urban context, for example. Importantly, the shared component of explained variance between urban context and land use was consistently minimal (<3%), indicating that these two variables act largely independently in shaping community composition. Notably, ESV-level patterns closely mirrored those at higher taxonomic ranks, supporting their use as sensitive and reliable units for ecological inference. Aligned with these results, pairwise comparisons revealed that agricultural versus forested sites consistently showed the most significant contrasts in community composition (median R² = 0.08-0.11 across amplicons), followed by weaker but significant differences between mosaic and agricultural or mosaic and forested sites (median R² = 0.045-0.078; Table 4). These results indicate that agricultural land use exhibits the strongest signal on community composition, whereas mosaic landscapes occupy an intermediate position between disturbed and less-impacted sites.

**Figure 6:**
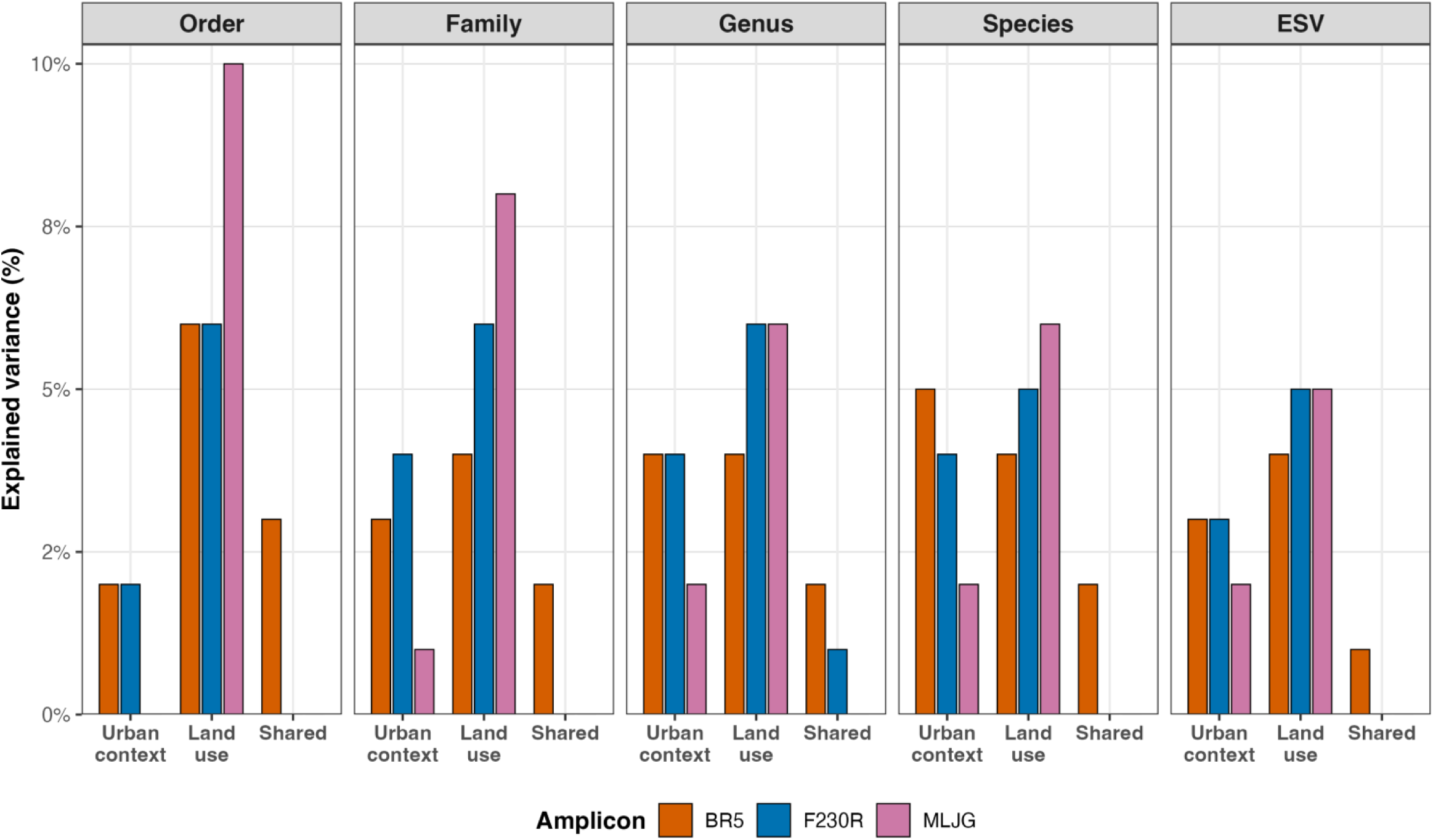
Partitioning of explained variance in DNA metabarcoding community composition across three COI amplicons (BR5, F230R, MLJG). Results are presented for presence/absence community matrices at all taxonomic levels investigated (ESV, species, genus, family, and order). Bars show the percentage of community variation explained by urban context, land use, and their shared variance, based on variance partitioning of PERMANOVA models. For all taxonomic levels, land use explained most of the variance across all amplicons.

**Table 4:** Summary of pairwise PERMANOVA results comparing macroinvertebrate community composition among land-use categories for each amplicon.

| Amplicon | Contrast | Levels | $R^2$ median<br>[min–max] | $p$ <sup>adj</sup> (min–max) |
| --- | --- | --- | --- | --- |
| BR5 | Predominantly Agricultural vs Predominantly Forested | ESV, Species, Genus, Family, Order | 0.109<br>[0.093–0.187] | 0.0009–0.001 ** |
| BR5 | Mosaic Landscape vs Predominantly Forested | ESV, Species, Genus, Family, Order | 0.063<br>[0.055–0.065] | 0.001–0.036 * |
| F230R | Predominantly Agricultural vs Predominantly Forested | ESV, Species, Genus, Family, Order | 0.093<br>[0.090–0.116] | 0.0009–0.012 * |
| F230R | Mosaic Landscape vs Predominantly Agricultural | ESV, Species, Genus, Family, Order | 0.078<br>[0.066–0.092] | 0.002–0.012 * |
| F230R | Mosaic Landscape vs Predominantly Forested | ESV, Species, Genus | 0.048<br>[0.048–0.049] | 0.013–0.030 * |
| MLJG | Predominantly Agricultural vs Predominantly Forested | ESV, Species, Genus, Family, Order | 0.080<br>[0.079–0.084] | 0.002–0.036 * |
| MLJG | Mosaic Landscape<br>vs Predominantly<br>Agricultural | ESV, Species,<br>Genus | 0.050<br>[0.049–0.053] | 0.030–0.043 * |
| MLJG | Mosaic Landscape<br>vs Predominantly<br>Forested | ESV, Species | 0.045<br>[0.045–0.046] | 0.030–0.036 * |

Only contrasts with significant differences at one or more taxonomic levels are shown. For each contrast, all significant taxonomic levels (FDR-adjusted $p < 0.05$ ) are listed in the “Levels” column. Effect sizes are shown as median $R^2$ values with their range across levels in brackets. Adjusted $p$ -values are reported as the minimum–maximum range across levels, with significance based on the highest (most conservative) $p$ -value per contrast. Significance levels are indicated as: $p < 0.05$ (\*), $p < 0.01$ (\*\*).

## Discussion

### DNA metabarcoding recovers more of the species pool

Our results indicate that, relative to the historical morphology dataset, DNA metabarcoding recovered substantially more orders, families, genera, and species. This pattern aligns with prior studies demonstrating that DNA metabarcoding often yields higher taxonomic coverage and finer resolution, by not just increasing richness estimates but capturing cryptic, soft-bodied, or otherwise overlooked taxa of ecological importance (Emilson et al., 2017; Pereira-da-Conceicoa et al., 2021). However, to our knowledge, our study is the first to provide a comparison of DNA metabarcoding from one year of sampling to morphological data captured in a span of 17 years. As such, our study shows how the increased information content can impact the outcome of the ecological analysis. From a methodological standpoint, DNA metabarcoding integrates signal from bulk tissue and trace DNA. The employment of multi-primer/marker assays (BR5, F230R, MLJG) with broad coverage for the COI gene reduces taxon-specific primer bias and increases the probability of detecting phylogenetically diverse lineages (Hajibabaei et al., 2012; Gibson et al., 2014; Elbrecht et al., 2017; Vamos et al., 2017; Hajibabaei et al., 2019).

These findings illustrate a well-known contrast between the two approaches: while morphology-based identifications captured the major macroinvertebrate groups, they were often restricted to a few abundant orders (e.g., Diptera, Coleoptera, and Trichoptera) and included thousands of unresolved taxa, whereas DNA metabarcoding provided broader taxonomic coverage and better resolution across multiple aquatic invertebrate lineages. It has been shown that morphology-based analysis is also biased towards taxa with larger body size, hence reducing the probability of documenting smaller organisms (Orlofske and Baird, 2013).

From a conservation and ecological management perspective, the fact that thousands of specimens remained unresolved at the genus/species level imposes a need for higher-yield methodologies that reduce collection pressure while improving taxonomic resolution. DNA metabarcoding of bulk samples has repeatedly shown the capacity to deliver high throughput, taxonomic breadth, and resolution suitable for teasing apart fine-scale changes in the communities (Hajibabaei et al., 2011; Elbrecht and Leese, 2017; Aylagas et al., 2018; Robinson et al., 2022).

### Spatial and environmental interactions in community structure

The observed structuring of benthic macroinvertebrate communities by urban development context likely reflects ecological gradients driven by differences in landscape composition. The urbanization context classification separates “large city” regions, characterized by dense urban development and adjacent agriculture, from “medium city” regions, where forested and transitional land uses predominate. Such alignment between urban context and land use is well-documented in watershed ecology, where spatial units (e.g., catchments, sub-watersheds, or administrative regions) frequently encompass broader land-use gradients that influence stream ecosystems (Allan, 2004). This overlap suggests that urban context-level groupings may integrate both spatial and environmental variation in community structure. Additionally, our results demonstrate that DNA metabarcoding not only distinguishes community differences among land-use types but also effectively captures their associations with environmental gradients at the genetic level. Prior work on a smaller scale supports this observation (Robinson et al., 2022). Variables potentially linked to agricultural activities, such as conductivity, pH, and turbidity, were consistently correlated with community composition across ESVs. Additionally, oxygen-related variables highlighted the ecological distinctiveness of forested sites, consistent with the expected cooler, shaded conditions of forested streams, which often maintain higher oxygen availability (Allan, 2004; Sweeney and Newbold, 2014; dos Reis Oliveira et al., 2019). Water turbidity was also a significant environmental variable, showing an expected outcome in tile-drained agricultural landscapes where two opposing processes may occur: (1) reduced riparian cover and soil disturbance promote higher runoff and sediment transport, while (2) tile drainage may locally dilute turbidity by introducing clearer groundwater into stream channels (Sweeney et al., 2004; Blann et al., 2009; Grangeon et al., 2021). Therefore, along with the statistical results, our findings point to associations between benthic community structure and broader environmental gradients such as water chemistry, geology, and local land-use patterns in the South Nation watershed. Similar trends have been reported before, where catchment-scale land use is strongly associated with stream condition, although the causal relationships remain complex and context-dependent (Roth et al., 1996; Morley and Karr, 2002; Allan, 2004; dos Reis Oliveira et al., 2019).

### Limited but significant variance in **β**-diversity metrics

In accordance with previous biomonitoring studies, we observed modest relationships between β-diversity and environmental gradients (Heino et al., 2013; Astorga et al., 2014; Keke et al., 2021). However, it is known that even in heterogeneous systems with carefully measured covariates, species-specific responses, dispersal/mass effects, and stochasticity can decrease community-environment correlations (Heino, 2015). Accordingly, the low but consistent effect sizes we report (PERMANOVA R² ≈ 0.08–0.11 for agricultural vs. forested contrasts) are typical for complex, species-rich freshwater assemblages rather than evidence of weak signals. Moreover, we show that urban context also acts as an ecological driver for macroinvertebrate communities in the south Nation watershed. Therefore, we argue for study designs that consider both spatial structure and land cover when interpreting multivariate biological community patterns. Small yet consistent explained variance is ecologically meaningful, especially when replicated across amplicons and taxonomic levels. Future work should (i) resolve indicator taxa for each land-use type to sharpen diagnostics and (ii) link community turnover to functional responses (e.g., transcriptomics) to clarify how physicochemical drivers translate into biological mechanisms.

### From complementarity to adoption: Why DNA metabarcoding belongs in routine biomonitoring

For years, studies have highlighted the importance of combining DNA metabarcoding with traditional morphological assessments for biomonitoring purposes (Hajibabaei et al., 2011; Baird and Hajibabaei, 2012; Elbrecht et al., 2017). However, concerns have been raised regarding the limitations of DNA-based methods, specifically their inability to provide quantitative estimates of abundance (Elbrecht and Leese, 2015; Lamb et al., 2019). The results from this study clearly indicate that DNA metabarcoding alone generates stronger and more consistent ecological signals compared to morphology-based approaches, particularly when detecting community-level responses to anthropogenic stressors, such as agricultural intensification and land-use changes. Without the fine-scale resolution offered by DNA metabarcoding, minimal but critical shifts in community structure may remain undetected, potentially compromising the effectiveness of management strategies and conservation decision-making (Pawlowski et al., 2018; Bush et al., 2019). Our comparisons further emphasize these differences: while the morphology dataset spanned more than a decade (2008-2022), its discriminatory power was limited, with some statistical results being weak or inconsistent across taxonomic levels. Group separation in morphology-based NMDS ordinations was often distorted by high within-group variability, particularly at family and genus levels, a result partially attributable to the high proportion of “Unidentified” specimens in the dataset, a clear methodological constraint. Another important consideration points to taxonomic expertise, which can introduce inconsistencies in detection and classification. These taxonomic gaps not only reduced resolution but also introduced artificial variation that may have interfered with the statistical results.

By contrast, although restricted to a one-year sampling period, DNA metabarcoding analyses consistently produced robust signals across all three amplicons and at finer taxonomic resolutions (ESVs, genus, and species). Its ability to capture cryptic, rare, and early instar taxa, alongside higher completeness of fine-rank data, greatly enhanced its statistical power. Moreover, variance partitioning confirmed that urban context and land-use effects were independently recovered across amplicons and taxonomic levels, with ESV-level patterns mirroring higher ranks. In general, our results also show that taxonomic resolution and detection sensitivity may be more critical than temporal scope in revealing ecological gradients.

Traditional morphological approaches remain useful, especially for trait-based inference and historical comparisons (Haase et al., 2010; Sweeney et al., 2011). Nevertheless, given the demonstrated robustness and reproducibility of DNA metabarcoding across multiple primer sets (BR5, F230R, MLJG) and taxonomic levels (ESV to Order), we strongly recommend the adoption of DNA metabarcoding into long-term biomonitoring programs, especially in sites that are highly impacted by mixed land-use stressors such as the South Nation watershed. Integrating DNA-based approaches in environmental assessment and monitoring will significantly enhance the ability to detect, monitor, and manage ecological changes in freshwater ecosystems (Baird and Hajibabaei, 2012; Leese et al., 2016; Deiner et al., 2017). The future of freshwater biomonitoring relies on an integrative framework in which DNA metabarcoding and other eDNA-based methodologies will provide a rapid first-pass analysis at a large scale, and more traditional surveys can provide complementary or confirmatory data where needed. Our study clearly demonstrates that a sensitive, timely, scalable, and comprehensive analysis of sites under land use change can be achieved even with a single sampling campaign, paving the way for better conservation and management in the face of accelerating environmental change.

## Acknowledgements

We would like to acknowledge the South Nation Conservation Authority (SNCA) for providing part of the data used in this paper, and Victoria Carley Maitland, Alysha Wenghofer, and Tessa Wynn for helping during the collection of the eDNA samples.

