## Supplementary material for "Fine-scale ecological biomonitoring in a large, complex agriculturally impacted watershed via eDNA metabarcoding": Tables S1, S3 and Figure S1

| Sampling year | Total specimens  collected | Unidentified Families | Unidentified Genus | Unidentified Species |
| --- | --- | --- | --- | --- |
| 2008 | 1283 | 132 (10.29%) | 1189 (92.67%) | 1283 (100%) |
| 2009 | 228 | 7 (3.07%) | 227 (99.56%) | 228 (100%) |
| 2010 | 309 | 6 (1.94%) | 309 (100%) | 309 (100%) |
| 2013 | 9003 | 90 (1.0%) | 8935 (99.24%) | 900 (100%) |
| 2017 | 2613 | 0 | 291 (11.14%) | 2424 (92.77%) |
| 2018 | 1916 | 178 (9.29%) | 273 (14.25%) | 1752 (91.44%) |
| 2019 | 29023 | 0 | 3338 (11.5%) | 25217 (86.89%) |
| 2020 | 14350 | 0 | 2709 (18.88%) | 11383 (79.32%) |
| 2021 | 7034 | 0 | 658 (9.35%) | 5796 (82.4%) |
| 2022 | 13538 | 0 | 1274 (9.41%) | 10546 (77.9%) |

**Table S1: Total number of arthropod specimens collected annually by South Nation Conservation under the Ontario Benthos Biomonitoring Network (OBBN) from 2008 to 2022, with the respective number and percentage of individuals recorded as “Unidentified” at the family, genus, and species levels.** While family-level assignments were rarely unresolved, a high proportion of specimens remained unidentified at the genus and especially at the species level, highlighting limitations of morphology-based identification.

| Variable | R² | R² significance | P value | Significance | Amplicon |
| --- | --- | --- | --- | --- | --- |
| Water temperature (°C) | 0.138 | Moderate | 0.047 | * | BR5 |
| pH | 0.215 | Moderate | 0.006 | ** | BR5 |
| Specific conductivity (µS/cm) | 0.534 | Very Strong | 0.001 | *** | BR5 |
| Dissolved O_2_ | 0.166 | Moderate | 0.032 | * | BR5 |
| Turbidity (NTU) | 0.116 | Moderate | 0.073 |  | BR5 |
| Dissolved O_2_ (%) | 0.078 | Weak | 0.179 |  | BR5 |
| ORP (mV) | 0.099 | Weak | 0.123 |  | BR5 |
| Pressure (mmHg) | 0.122 | Moderate | 0.048 | * | BR5 |
| Water temperature (°C) | 0.333 | Strong | 0.001 | *** | F230R |
| pH | 0.273 | Moderate | 0.002 | ** | F230R |
| Specific conductivity (µS/cm) | 0.434 | Strong | 0.001 | *** | F230R |
| Dissolved O_2_ | 0.127 | Moderate | 0.073 |  | F230R |
| Turbidity (NTU) | 0.150 | Moderate | 0.043 | * | F230R |
| Dissolved O_2_ (%) | 0.037 | Very Weak | 0.462 |  | F230R |
| ORP (mV) | 0.021 | Very Weak | 0.634 |  | F230R |
| Pressure (mmHg) | 0.101 | Moderate | 0.129 |  | F230R |
| Water temperature (°C) | 0.293 | Moderate | 0.001 | *** | MLJG |
| pH | 0.375 | Strong | 0.001 | *** | MLJG |
| Specific conductivity (µS/cm) | 0.504 | Very Strong | 0.001 | *** | MLJG |
| Dissolved O_2_ | 0.100 | Moderate | 0.103 |  | MLJG |
| Turbidity (NTU) | 0.176 | Moderate | 0.024 | * | MLJG |
| Dissolved O_2_ (%) | 0.081 | Weak | 0.171 |  | MLJG |
| ORP (mV) | 0.005 | Very Weak | 0.884 |  | MLJG |
| Pressure (mmHg) | 0.253 | Moderate | 0.003 | ** | MLJG |

**Table S3: Results of envfit analyses to test the relationship between environmental variables and community composition for three COI amplicons (BR5, F230R, MLJG).** Reported values include the coefficient of determination (R²), qualitative interpretation of effect size (very weak to very strong), and permutation p-values (999 iterations). Significance codes: p < 0.05 (*), p < 0.01 (**), p < 0.001 (***).

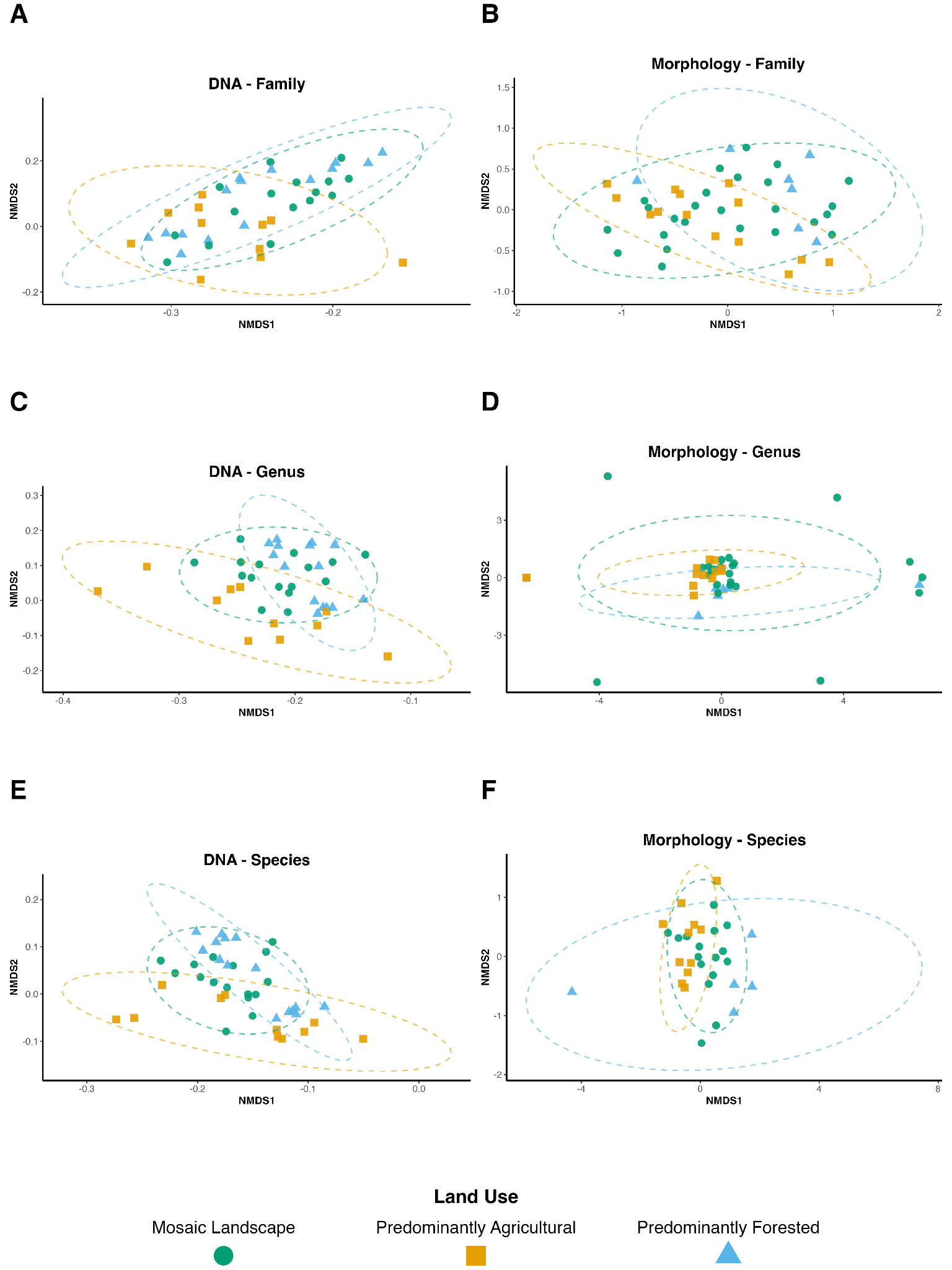

**Figure S1: Non-metric multidimensional scaling (NMDS) ordination plots showing macroinvertebrate community composition across land use types using DNA metabarcoding (left column) and morphology-based identification (right column) at three taxonomic levels: family (A, B), genus (C, D), and species (E, F).** Ordinations were computed using Bray-Curtis dissimilarities, based on presence/absence data for DNA (F230R amplicons) and abundance data for morphology. DNA results are based only on one year of sampling (2023), while morphology results represent ten years of sampling. Points represent individual samples, and 95% confidence ellipses group sites by land use category: mosaic landscape (green circle), predominantly agricultural (orange square), and predominantly forested (blue triangle). All morphology-based samples are shown, including outliers. Stress values and ordination: DNA - family (0.148, k=2), DNA - genus (0.155, k=2), DNA - species (0.146, k=2); Morphology - family (0.164, k=3), Morphology - genus (0.067, k=2), Morphology - species (0.187, k=2).
